# A Glu-Glu-Tyr sequence in the cytoplasmic tail of the M2 protein renders IAV susceptible to restriction of HA-M2 association in primary human macrophages

**DOI:** 10.1101/2020.07.01.183186

**Authors:** Sukhmani Bedi, Amanda Haag, Akira Ono

## Abstract

Influenza A virus (IAV) assembly at the plasma membrane is orchestrated by at least five viral components including hemagglutinin (HA), neuraminidase (NA), matrix (M1), the ion channel M2, and viral ribonucleoprotein (vRNP) complexes although particle formation itself requires only HA and/or NA. While these five viral components are expressed efficiently in primary human monocyte-derived macrophages (MDM) upon IAV infection, this cell type does not support efficient HA-M2 association and IAV particle assembly at the plasma membrane. The defects in HA-M2 association and particle assembly are specific to MDM and not observed in a monocytic cell line differentiated into macrophage-like cells. Notably, both these defects can be reversed upon disruption of the actin cytoskeleton. In the current study, we sought to examine whether M2 contributes to particle assembly in MDM and to identify a viral determinant involved in the MDM-specific and actin-dependent suppression of IAV assembly. An analysis using correlative fluorescence and scanning electron microscopy showed that an M2-deficient virus fails to form budding structures at the cell surface even after F-actin is disrupted, indicating that M2 is essential for virus particle formation at the MDM surface. Notably, proximity ligation analysis revealed that single amino acid substitution in a Glu-Glu-Tyr sequence (residues 74-76) in the M2 cytoplasmic tail allows HA-M2 association to occur efficiently even in MDM with intact actin cytoskeleton. This phenotype did not correlate with known phenotypes of the M2 substitution mutants regarding M1 interaction or vRNP packaging in epithelial cells. Overall, our study identifies a viral determinant for susceptibility to cytoskeleton-dependent regulation in MDM and hence, sheds light on the molecular mechanism behind the MDM-specific restriction of IAV assembly.

**Importance:** Non-permissive cell types that are unable to support viral replication serve as important tools for identification of host factors that either block viral replication (restriction factors) or support viral replication in permissive cell types (co-factors). We previously identified the MDM as a cell type that is non-permissive to IAV assembly, likely due to a block in HA-M2 association. In the current study, we determined that the IAV M2 protein is necessary for virus particle formation in MDM but also renders the virus susceptible to the MDM-specific suppression of virus assembly. We identified a specific amino acid motif in the M2 cytoplasmic tail, disruption of which allows M2 to associate with HA even in MDM. Our findings strongly support the possibility that the MDM-specific defect in HA-M2 association is due to the presence of a restriction factor(s) in MDM, which likely interacts directly with the M2 cytoplasmic domain, rather than indirectly through other internal viral components, and thereby prevents M2 from associating with HA.

## Introduction

IAV buds from areas enriched in cholesterol-rich microdomains or lipid rafts, which are also known as budozones, of the plasma membrane (1–3). IAV transmembrane proteins HA and NA intrinsically localize to lipid rafts of the apical plasma membrane (4–8), which eventually coalesce into larger domains (5, 6, 8–10). HA and/or NA expression at the plasma membrane has been shown to be sufficient for induction of curvature for spherical virus bud formation (11–13). The third transmembrane protein M2 also intrinsically traffics to the apical plasma membrane (14); however, it does not associate with lipid rafts when expressed by itself (4). In infected cells, M2 is shown to associate closely with HA (15, 16) while still localizing to the boundary of lipid rafts (5, 17–20). In contrast with HA and NA, M2 is thought to be dispensable for curvature induction but rather plays important roles in pinching off of the spherical virus bud prior to virus release (17, 19–21).

In addition to the role of M2 in pinching off of the virus bud prior to virus release, multiple lines of evidence suggest that M2 also plays a role(s) during the earlier stages of IAV particle assembly. First, deletion of M2 (18) or introduction of mutations in M2 (19, 20, 22–24) alter the morphology of assembling virus particles. Second, incorporation of M1 into assembling virus particles, which is important for providing structure to the virus particle, is at least partially dependent on M2 (24–26). The cytoplasmic tail of the M2, more specifically amino acid residues 71-78, has been implicated as an interface for M2-M1 association (25, 26). Third, the incorporation of vRNP into the assembling virus particle, which further promotes IAV assembly (27, 28), is also at least partially dependent on M2 (22–24, 26, 29–31). As for the molecular determinants, some of the same amino acid residues implicated in incorporation of M1are shown to be important for incorporation of vRNP into the virus particle (22–24, 26, 29, 30). Despite these lines of evidence, the exact role of M2 early during the IAV assembly process remains to be determined.

Previously, we characterized the primary human monocyte-derived macrophage (MDM) as a cell type that is inefficient at IAV particle assembly/release (15). We found that HA-M2 association, defined as localization of the two proteins in a close (<40 nm) proximity of each other, is diminished in this cell type. In contrast, macrophage-like cells derived from the THP1 human monocytic cell line support both efficient HA-M2 association and virus particle assembly. The MDM-specific defect in HA-M2 association is reversed upon disruption of the actin cytoskeleton, suggesting the presence of an Factin-dependent host regulation of HA-M2 association. Importantly, we further found that actin disruption in MDM restores virus particle formation detected as budding structures on the cell surface, suggesting a strong correlation between M2 recruitment to the proximity of HA and virus assembly/budding in this cell type.

In this study, we elucidate the molecular determinant essential for the F-actin-dependent suppression of IAV assembly in MDM. Using an M2-deficient virus, we show that the expression of M2 is required for efficient particle assembly in MDM upon Factin disruption, a condition under which IAV assembly is supported in this cell type. Consistent with an important role for M2 in IAV assembly in MDM, efficient IAV assembly in differentiated THP1 (dTHP1) cells, which are permissive to IAV assembly, is also dependent on the expression of M2. Hence, in addition to its previously described role in scission of the virus bud late during IAV assembly in epithelial cells, M2 plays a role(s) early during virus particle formation in macrophages, which may overlap with the roles M2 play in particle morphology or vRNP incorporation (18–20, 22–26, 29–31). Furthermore, we found that substitutions of three individual amino acids in the cytoplasmic tail of M2 (E74, E75, and Y76), but not the preceding three residues, allow efficient HA-M2 association in MDM, suggesting the presence of a restriction factor that recognizes this sequence. Altogether, our data indicate that M2 is necessary for formation of IAV particles at the MDM surface but it also confers to IAV the susceptibility to a cell-type-specific restriction by the actin cytoskeleton via its cytoplasmic tail.

## Materials and methods

### Cells and reagents

Monocytes were isolated by plate adhesion from peripheral blood mononuclear cells, which were obtained from buffy coats derived from unidentified healthy donors (New York Blood Center, NY). Cells were cultured in RPMI 1640(Gibco) supplemented with 10% fetal bovine serum (FBS, Hyclone) for 7 days before they were used for experiments. THP1 (ATCC® TIB202™) cells were cultured in RPMI 1640 supplemented with 10% FBS, 1 mM Sodium Pyruvate (Gibco) and 0.05 mM 2-mercaptoethanol. To generate differentiated THP1 cells (dTHP1), THP1 cells were cultured in the medium containing 0.1 μM phorbol 12-myristate 13-acetate (PMA; Sigma) and 0.1 μM Vitamin D3 (Sigma) for 2-3 days. Madin-Darby canine kidney (MDCK) cells were provided by Dr. Arnold S. Monto (University of Michigan) and were cultured in DMEM (Gibco) supplemented with 10% FBS and 25 mM HEPES. Human embryonic kidney-derived 293T cell line (ATCC) was cultured and maintained in DMEM (Lonza) supplemented with 10% FBS.

The following antibodies were used for immunofluorescence microscopy and/or flow cytometry: mouse anti-HA monoclonal antibody (clone C179 (32); Takara), mouse anti-M2 monoclonal antibody (clone 14C2 (33); Thermofisher), goat anti-HA antiserum (BEI NR-3148), rabbit anti-M1 (GTX125928; GeneTex). All secondary antibodies used for immunofluorescence were purchased from Thermofisher. Latrunculin B was purchased from Sigma and re-constituted in DMSO.

### Generation of M2-MDCK stable cell line

cDNA corresponding to the M2 protein was cloned into the pCABSD plasmid (a kind gift from Dr. F. Momose), which expresses a blasticidin S resistance gene. MDCK cells were transfected with the pCABSD-M2 plasmid using lipofectamine (Invitrogen). Cells stably expressing M2 were selected in growth medium containing 1 μg/ml blasticidin and were enriched for high M2 expression by staining with an anti-M2 antibody (clone 14C2) and an Alexa 488-conjugated secondary antibody followed by fluorescence-activated cell sorting (FACSAria II, BD Biosciences).

### Generation of plasmids encoding M2 mutants

Generation of M2 mRNA from the WSN M segment was disrupted as described previously (34). Briefly, a WSN M segment with a disrupted splicing signal (WSN M SS) was generated by introducing mutations in the 3’ splice site of the pPOLI-WSN-M plasmid using QuikChange II site-directed mutagenesis protocol (Agilent). pPolI-WSN-M-M2SMRtoAAA (Mut1), pPolI-WSN M-M2EEYtoAAA (Mut2), pPolI-WSN M-M2E74A, pPolI-WSN M-M2E74A, and pPolI-WSN M-M2Y76A constructs were also generated using QuikChange II site-directed mutagenesis protocol.

### Generation of virus stocks

A/WSN/1933 (H1N1) virus was generated by reverse genetics (35) using the 8 pPolI plasmids encoding different segments of IAV genome and the 4 pCAGGS plasmids that express the PA, PB1, PB2, and NP proteins. ΔM2 and M2 mutant WSN viruses were generated in 293T cells using the pPolI plasmids encoding the 7 RNA segments of the WT WSN strain, pPolI plasmid encoding mutant WSN M2 segment, and the 5 pCAGGS plasmids that express the PA, PB1, PB2, NP, and M2 proteins. Virus generated from 293T cells was propagated in M2-MDCK cells. Viral titers were determined by plaque assays performed using M2-MDCK cells.

### Virus infection

For virus infection, cells were first washed twice with MEM-BSA medium (1X MEM supplemented with 0.3% BSA (Sigma), 0.23% sodium bicarbonate (Gibco), 1X MEM amino acids (Gibco), and 1X MEM vitamins (Gibco)). The cells were then inoculated with diluted virus at MOI 0.01 or 0.1 for 1 hour at 37°C, washed twice with MEM-BSA, and cultured in MEM-BSA containing 0.2 μg/ml tosylsulfonyl phenylalanyl chloromethyl ketone (TPCK)-treated trypsin (Worthington).

### Flow cytometry

For viral protein expression analysis by flow cytometry, virus- or mock-infected cells were detached using 0.06% trypsin-EDTA in phosphate-buffered saline (PBS) and fixed with 4% paraformaldehyde (PFA, Electron Microscopy Sciences) in PBS for 20 minutes at room temperature. After fixation, cells were washed twice with 2% FBS in PBS (FACS buffer). For detecting viral protein expression on the cell surface, cells were subsequently incubated with primary antibodies for 45-60 minutes directly. For staining of intracellular proteins, cells were first permeabilized with 0.1% TritonX-100 in PBS for 5 minutes and then probed with primary antibodies. After washing once with the FACS buffer, cells were incubated with fluorescently labeled secondary antibodies (Invitrogen) for 20-30 minutes. Cells were washed twice with the FACS buffer and analyzed using the FACSCanto flow cytometer (BD Biosciences). Data were analyzed in FlowJo (Treestar), and the positive gates were set using the mock-infected controls.

### Correlative fluorescence and scanning electron microscopy (CFSEM)

CFSEM experiments were performed as described before (15, 36). Briefly, cells cultured on gridded coverslips (Bellco Biotechnology) were infected with WSN at MOI 0.1. Cells were fixed with 4% PFA in PBS at 20 hpi. After rinsing in PBS, quenching of PFA with PBS containing 0.1 M glycine (Sigma), and blocking with PBS containing 3% bovine serum albumin (BSA, Sigma), cells were immunostained with mouse anti-HA and fluorescently labeled secondary antibody. Cells were imaged using a Leica Inverted SP5X Confocal Microscope with a 40× PL APO objective and 10-20× scanning zoom. After fluorescence imaging, cells were fixed with PBS containing 2.5% glutaraldehyde (Electron Microscopy Sciences), stained with 1% OsO4, dehydrated in a series of ethanol washes, rinsed in hexamethyldisilazane (Electron Microscopy Sciences) and allowed to dry overnight. Coverslips were affixed to specimen mounts and sputter coated with gold for 90 s (Polaron). Cells were identified by their location on the gridded coverslip and imaged on an Amray 1910FEG scanning electron microscope at 5-10 kV. Fluorescence and SEM images were roughly brought into registration by scaling and rotating images in Adobe Photoshop, similarly to previous correlative fluorescence/SEM studies (15, 36). Landmarks used for registration included cell edges. Cell surface structures visible in SEM were manually classified as virus-like buds if they appeared spherical and near 100 nm in diameter. To identify HA clusters in fluorescence images unambiguously, we removed uniform non-puncta HA signal from the images. To do this, we calculated a 20-pixel radius median filter and subtracted the median filtered image from the original using the *Image Calculator* function in ImageJ. Since MDM have substantial membrane folds on the cell surface especially towards the center of the cell, we focused on areas towards the edge of the cells, which have a flatter topology, for quantification of efficiency of virus bud formation.

*In situ* Proximity Ligation Assay (PLA): PLA was performed using Duolink® PLA fluorescence kit (Sigma) as described previously (15). Cells fixed with 4% PFA (non-permeabilized) were incubated with goat anti-HA and mouse anti-M2 primary antibodies. Detection of PLA signals was performed using PLA probes specific to goat and mouse IgG. AlexaFluor-488-labeled secondary antibody was used to recognize anti-M2 and hence, to identify M2-positive cells. Cells were observed using a Leica Inverted SP5X Confocal Microscope System with a 63X objective. Z-stacks extending from the focal plane corresponding to the middle plane of the nucleus (identified by DAPI staining) to the bottom of cells were acquired for each cell, and the maximum intensity projection for each cell was constructed using ImageJ. The PLA signal in projection images was thresholded to eliminate weak and hazy background signal in the nucleus, and the number of PLA-positive spots was counted using the *Analyze particle* function in ImageJ. Statistical analysis: Statistical analyses were performed using GraphPad Prism version 8. Two-tailed paired student *t* test was used to calculate p-values in Figures 1 and 6. Twotailed unpaired student *t* test was performed in Figures 3–5.

## Results

### M2 is required for efficient IAV assembly in both MDM treated with Latrunculin B and dTHP1 cells

We previously showed that the actin cytoskeleton suppresses HA-M2 association and IAV assembly in MDM (15). However, it is not clear whether HA-M2 association precedes and is essential for assembly of IAV particles in MDM or whether IAV particle assembly is initiated independent of M2 upon F-actin disruption and HA-M2 association occurs post assembly of the virus particle. To address whether M2 is essential for IAV particle assembly in MDM, we generated a mutant virus (A/WSN/33 strain) that fails to express M2 (WSN ΔM2) in infected cells. This virus packages the WSN M segment with a disrupted 3’ splice site (WSN M SS), which allows for generation of the M1-encoding mRNA but not the M2-encoding mRNA from the M segment (18, 34) (Figure 1A). Since the stock WSN ΔM2 virus was prepared using MDCK cells stably expressing the WSN M2 protein, M2 is incorporated into virus particles and is able to complete its roles during the entry of IAV into cells; however, the virus is defective in de novo synthesis of M2, which is required during IAV assembly. To determine whether other viral structural proteins are expressed at comparable levels between WT virus- and ΔM2 virus-infected cells, we used flow cytometry to measure the total expression of M1 and cell surface expression of HA in MDM infected with WT or ΔM2 virus. For comparison with MDM, we used the monocytic cell line THP1, which has been differentiated to adopt macrophage-like morphology (dTHP1). dTHP1 cells, unlike MDM, support all steps of the IAV assembly process and have been used for comparison with MDM in our previous study (15). dTHP1 cells and MDM were infected with WT or ΔM2 virus at MOI 0.1 for 18 hours, after which cell surface expression of HA was measured in non-detergent-permeabilized cells, and total expression of M1 was measured in detergent-permeabilized cells. In addition, we measured the expression of M2 on the cell surface. As expected, dTHP1 cells and MDM infected with the ΔM2 virus failed to express M2 on the cell surface. Both HA and M1 were expressed in ΔM2 virus-infected MDM at levels comparable to that in WT virus-infected cells (Figure 1B). Cell surface expression of HA was about 2-fold reduced in dTHP1 cells infected with ΔM2 virus, relative to WT virus (Figure 1C). Total M1 expression was comparable between dTHP1 cells infected with WT and ΔM2 virus. Overall, our data indicate that in MDM, cell surface expression of HA and total expression of M1 are comparable between cells infected with ΔM2 and WT viruses.

**Figure 1:**
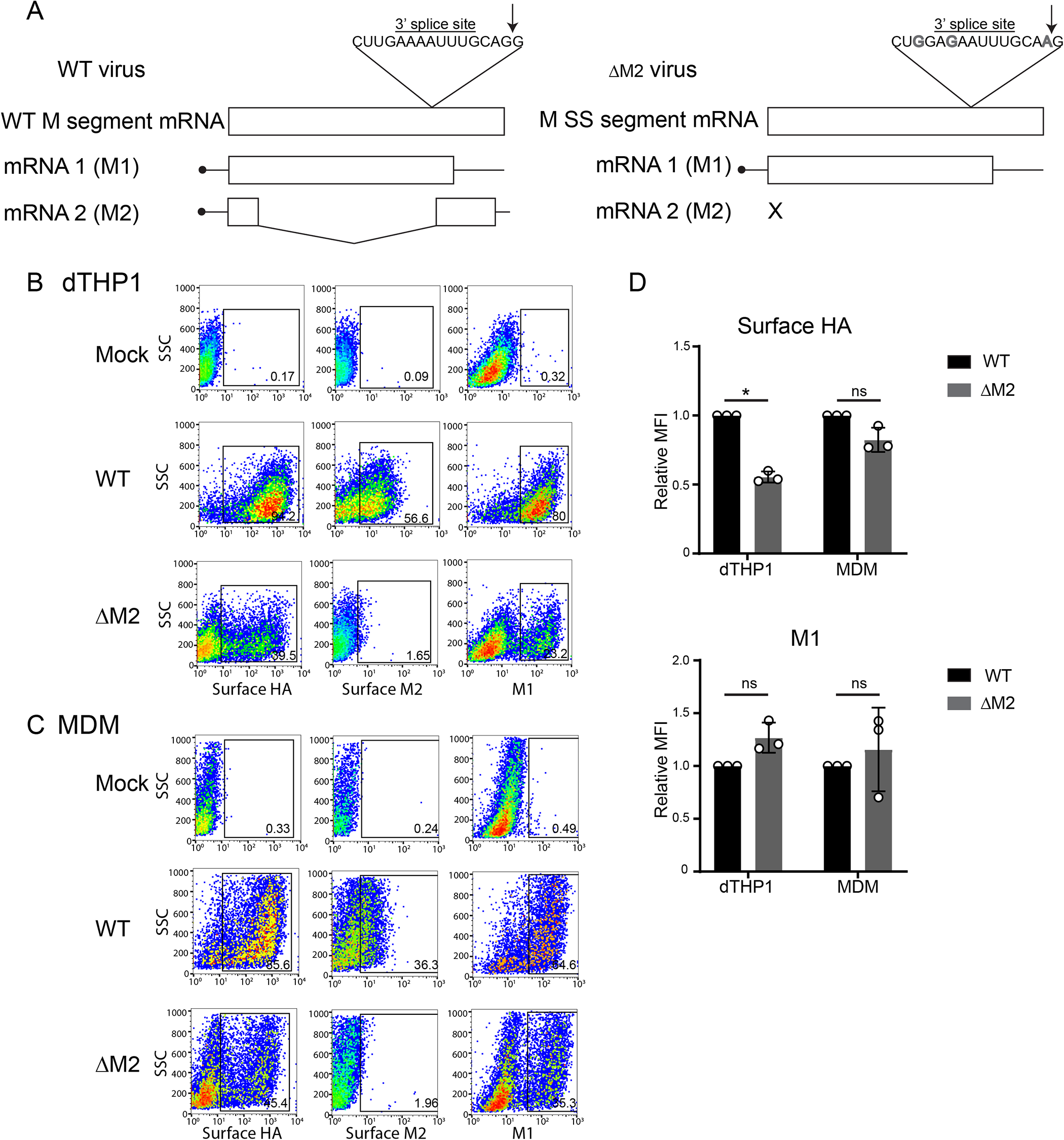
Generation and characterization of ΔM2 WSN virus. (A) The strategy used for generation of the WSN M segment that fails to encode for the M2 protein (WSN M SS) is depicted. (B-D) dTHP1 cells (B) and MDM (C) were infected with WT or ΔM2 virus at MOI 0.1 for 18 hours. (B and C) Infected cells were analyzed for cell surface expression of HA and M2 and intracellular expression of M1 by flow cytometry. Representative flow plots for mock-infected and virus-infected cells are shown. Percentages of cells positive for viral proteins (boxed) are shown. (D) Relative MFIs for signal of indicated proteins for positive cell populations (gated in panels B and C) are shown. Data are shown as mean +/-SD and are from at least three independent experiments. *, p<0.05; ns, non-significant.

We next asked whether restoration of IAV particle assembly upon F-actin disruption in MDM depends on expression of M2. To this end, we performed correlative fluorescence and scanning electron microscopy (CFSEM) with MDM infected with WT or ΔM2 virus and treated either with vehicle or 5 μM Latrunculin B (Lat B) at 14 hpi for 4 hours and examined virus bud formation in cells expressing HA on the cell surface. We counted the number of virus particle-like buds (~100 nm in diameter) within the same sized area (100 μm^2^ in size) of each cell. Consistent with previous results, WT virus-infected, vehicle-treated MDM showed very few buds on the cell surface. Similar to what was previously observed with cytochalasin D (15), disruption of the actin cytoskeleton with Lat B significantly increased the number of buds on the surface of WT virus-infected MDM (Figures 2A and 2B). As expected, very few bud-like structures were observed on the surface of vehicle-treated, ΔM2 virus-infected cells. Notably, no increase in number of buds was observed on the surface of ΔM2 virus-infected MDM upon treatment with Lat B, relative to treatment with vehicle (Figures 2A and 2B). These data indicate that disruption of the actin cytoskeleton promotes IAV particle assembly in MDM in an M2-dependent manner.

**Figure 2:**
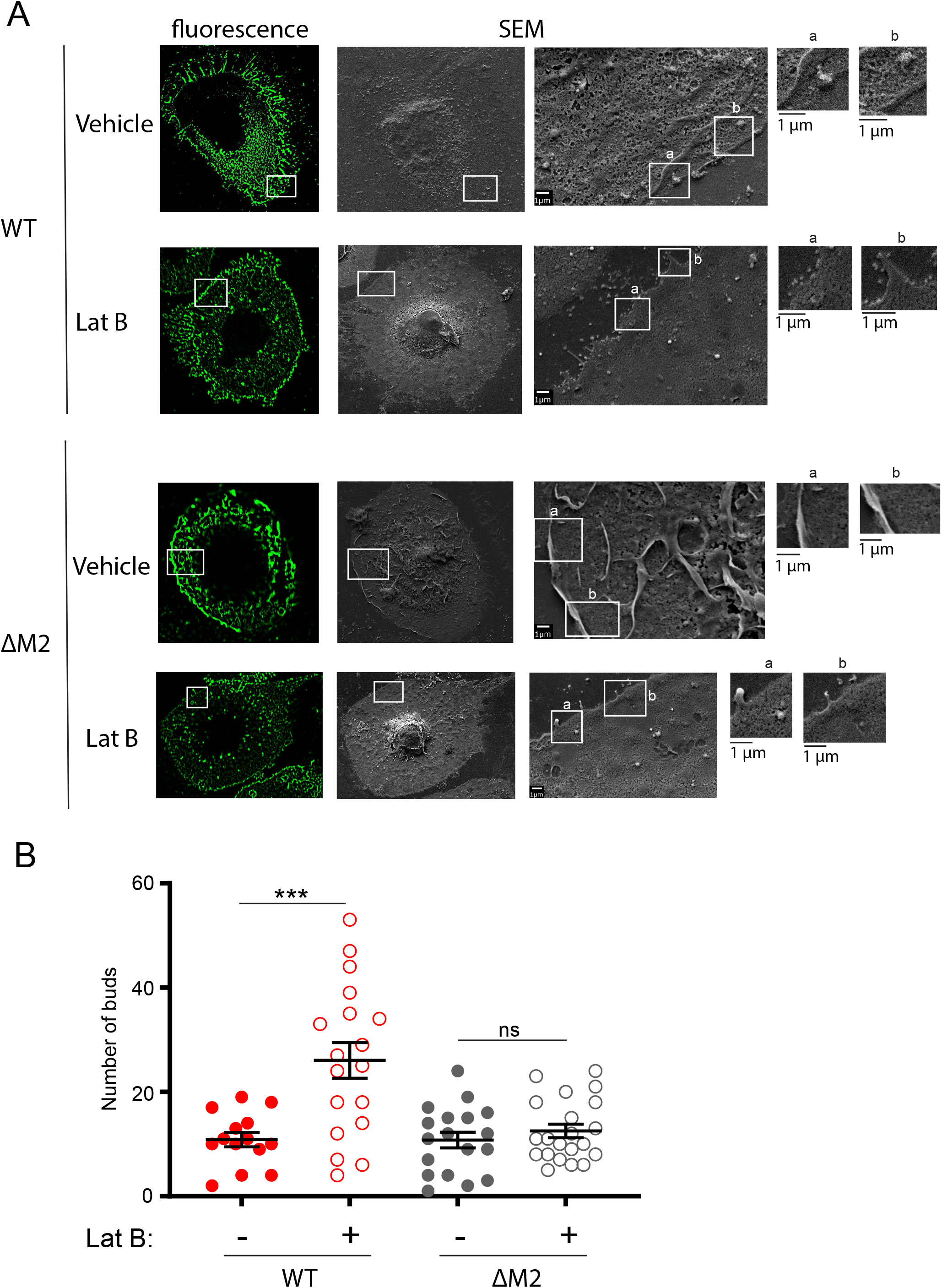
F-actin disruption restores virus bud formation in MDM in an M2-dependent manner. MDM were infected with WSN at MOI 0.1 for 14 hours. Infected cells were treated with vehicle (DMSO) or 5 μM Lat B for 4 hours, before fixation and immunostaining with anti-HA. After identification of HA-positive cells by confocal fluorescence microscopy, cells were processed for SEM. The same cells were identified based on grid positions and analyzed by SEM. (A) Representative SEM images for WSN-infected HA-positive cells are shown. Fluorescence images corresponding to the SEM images are also included. Boxed areas for SEM images are magnified and shown on the right of original images. Alphabetic labels are used to distinguish between the individual boxed areas. (B) The number of ~100-nm buds identified in SEM images were counted within the same sized area (100 μm^2^ in size) in each cell. Data are shown for 10-20 cells from two independent experiments. Error bars represent standard error of mean. ***, P<0.005; ns, non-significant.

Previous overexpression studies have shown that M2 is not essential for particle formation (11, 13). Consistent with this notion, IAV particle assembly is observed in an IAV-permissive epithelial cell line, MDCK, infected with virus that does not express M2 (17, 23). Therefore, we next asked whether dTHP1 cells, which are also permissive to IAV assembly, require M2 for particle formation on the cell surface. dTHP1 cells were infected with WT or ΔM2 virus and virus bud formation was analyzed on the surface of HA-positive cells by SEM. As expected, cells infected with the WT virus showed abundant bud-like structures on the cell surface. Notably, the number of buds observed on the surface of ΔM2 virus-infected dTHP1 cells was significantly reduced in comparison to that on WT virus-infected cells (Figures 3A and B). Since HA expression on the surface of ΔM2 virus-infected dTHP1 cells was reduced in comparison to WT virus-infected dTHP1 cells (Figure 1B), the reduction in bud formation in ΔM2 virus-infected cells may be due to the reduction in HA expression on these cells. However, in these experiments, cells expressing comparable levels of HA on the surface were analyzed for bud formation using SEM. This is more clearly observed upon measuring the fluorescence intensity of surface HA staining in individual dTHP1 cells, which were analyzed by SEM, using the ImageJ software. The fluorescence intensity of surface HA staining was comparable between the WT virus- and ΔM2 virus-infected dTHP1 cells analyzed by SEM (Figure 3C). Hence, the difference in surface HA expression does not explain the difference in bud formation between WT virus-versus ΔM2 virus-infected dTHP1 cells. These data indicate that efficient IAV particle formation relies on M2 in both dTHP1 cells and MDM. Overall, these data suggest that, in addition to its role in bud scission late during IAV assembly, M2 plays important roles early during IAV particle formation in MDM and dTHP1 cells.

**Figure 3:**
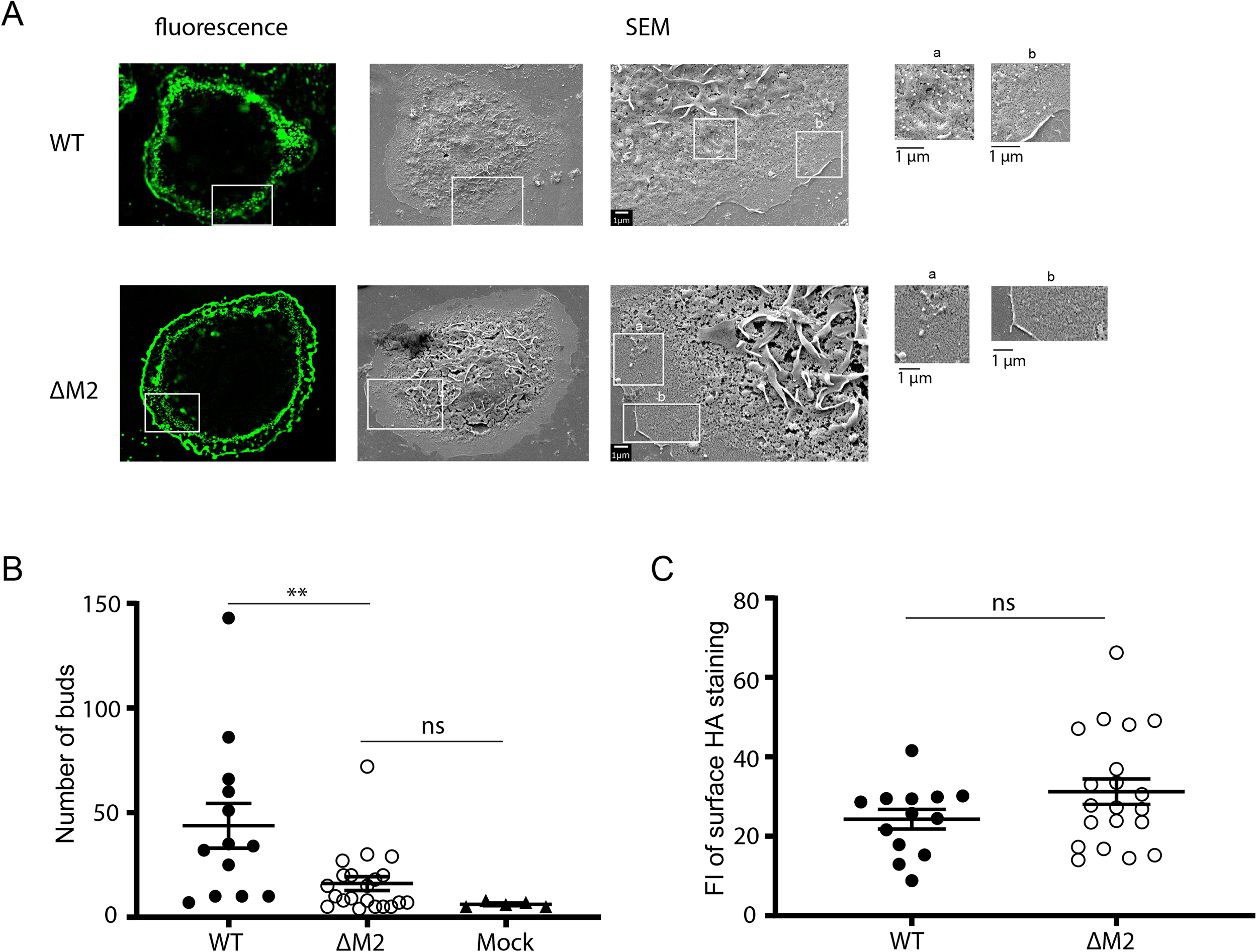
dTHP1 cells support IAV particle assembly in an M2-dependent manner. dTHP1 cells were infected with WT or ΔM2 virus for 18 hours. Infected cells were fixed and immunostained with anti-HA antibody. After identification of HA-positive cells by confocal fluorescence microscopy, cells were processed for SEM. The same cells were identified based on grid positions and analyzed by SEM. (A) Representative SEM images for WSN-infected HA-positive cells are shown. Fluorescence images corresponding to the SEM images are also included. Boxed areas for SEM images are magnified and shown on the right of original images. Alphabetic labels are used to distinguish between the individual boxed areas. (B) The number of ~100-nm buds identified in SEM images were counted within the same sized area (100 μm^2^ in size) in each cell. (C) Fluorescence intensity (FI) of surface HA expression was determined for each cell analyzed for virus bud formation using SEM. Data are shown for 10-20 cells from two independent experiments. Error bars represent standard error of mean. **, P<0.01; ns, non-significant.

### Amino acid residues 74-76 but not 71-73 in the cytoplasmic tail of M2 are important for the regulation of HA-M2 association in MDM

HA and NA associate efficiently on the surface of MDM but HA-M2 association is defective, suggesting that association of M2 with HA is a discrete step during IAV assembly that likely occurs independent of HA-NA association (15). The experiments above demonstrated that M2 is an important component that enables IAV particle assembly in MDM. Thus, the inefficient recruitment of M2 to the assembly sites enriched in HA is the likely cause of defective particle assembly in MDM. Since the defect in M2 association with HA is dependent on the intact actin cytoskeleton, we next sought to elucidate the viral motifs in M2 that are important for this F-actin-dependent regulation of HA-M2 association. At least two non-mutually exclusive mechanisms can be proposed for the MDM-specific F-actin-dependent restriction on M2: (i) an F-actin-dependent host factor directly interacts with M2 and limits its association with HA; (ii) an F-actin-dependent host factor interacts with a viral protein that associates with M2 to limit its association with HA (37).

To distinguish between these two proposed mechanisms, we focused on determining whether any sequence motifs on the M2 protein that have previously been implicated in mediating its associations with other viral proteins are also involved in regulating its association with HA. The two most studied M2-associating viral components are: M1 and vRNP (see Introduction). Notably, M1 (38) and NP (39) are also known to associate with F-actin and hence, could bridge interactions between M2 and Factin. Therefore, to determine whether association between M2 and M1 and/or vRNP is important for the restriction of HA-M2 association in MDM, we generated M2 constructs harboring mutations at amino acids 71-76, which have been previously shown to be important for association of M2 with M1 (25) and/or vRNP (22–24). We first tested two M2 mutants, with either amino acid residues 71-73 (Mut1) or 74-76 (Mut2) changed to alanine (25), for HA-M2 association using the previously described in situ Proximity Ligation Assay (PLA) (15) (Figure 4A). PLA allows for detection of two proteins localized within 40-nm distance of each other and has been used to visualize IAV assembly sites on the plasma membrane (15, 40). In addition to measuring PLA signal between HA and M2, we also co-stained cells for cell surface M2 to identify infected cells. As expected, very few PLA spots were observed between HA and M2 on MDM infected with the WT virus. Likewise, very few PLA spots were also observed between HA and Mut1 M2 (Figure 4B). Remarkably, in contrast with WT and Mut1 viruses, the number of PLA spots between HA and M2 increased drastically in Mut2 virus-infected MDM even without F-actin disruption (Figure 4B). Notably, no difference in the number of PLA spots was observed between WT and mutant viruses in infected dTHP1 cells (Figure 4C). This is unexpected, since Mut1 M2 has previously shown to be less efficient at co-clustering with HA at the plasma membrane of epithelial cells, relative to WT M2, in an electron microscopy-based study (16). However, this discrepancy between the previous study and our study could be attributed to differences in techniques (electron microscopy can detect a change in HA-M2 association at a shorter distance than PLA). Overall, our data suggest that amino acid residues 74-76 in the cytoplasmic tail of M2 regulate its association with HA in MDM. Of note, although substitutions in both Mut1 and Mut2 M2 are known to reduce interaction with M1 (25), they show different phenotypes in terms of restoring HA-M2 association in MDM. Hence, the defective association of M2 with HA in MDM does not correlate with the ability for M1-M2 association and therefore is unlikely to involve M2 association with M1.

**Figure 4:**
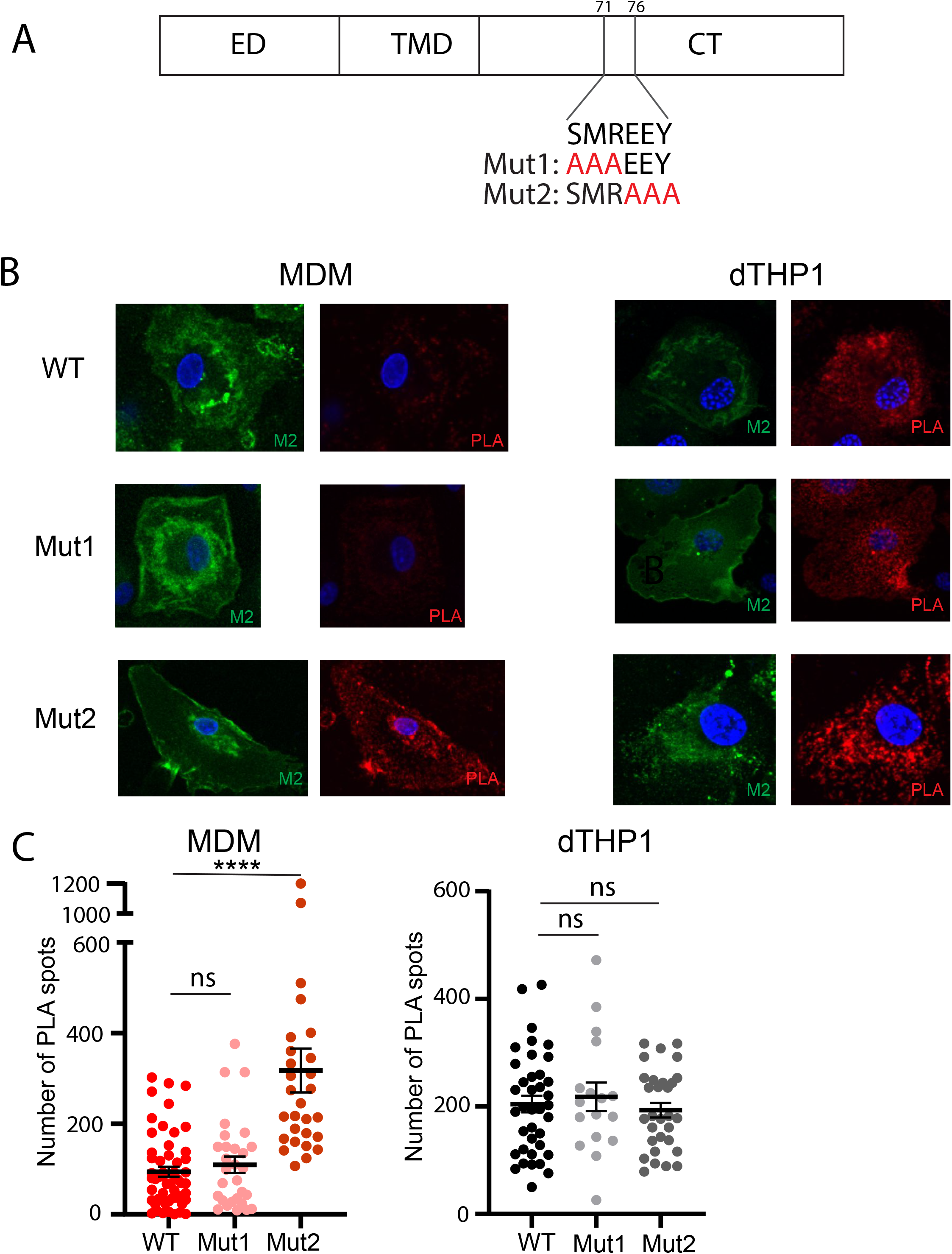
Alanine substitutions at amino acid positions 74-76 of the cytoplasmic tail of M2 restore HA-M2 association in MDM. (A) A schematic showing the mutations introduced in the cytoplasmic tail of M2. Amino acid residues 71-73 or 74-76 were substituted with alanines to generate Mut1 and Mut2 M2 mutants, respectively. (B, C) dTHP1 cells and MDM were infected with WT WSN or M2 mutant viruses at MOI 0.1 for 18 hours. Cells were fixed and examined by PLA using goat anti-HA and mouse antiM2 antibodies. To identify infected cells, total surface M2 was also detected using a fluorescently-tagged anti-mouse IgG secondary antibody (shown in green). Nuclei were stained with DAPI (blue). (B) Representative maximum intensity projection images, which were reconstructed from z-stacks corresponding to the focal planes ranging from the middle plane of the nucleus to the bottom of the cells, are shown. (C) Number of PLA spots were counted for each cell. Data are shown for experiments with MDM from three independent donors, and 8-10 cells were analyzed per experiment. Error bars represent standard error of mean. ****, p <0.0001; ns, non-significant.

### Amino acid residues at positions 74, 75, and 76 of the M2 protein are individually important for the defect in HA-M2 association in MDM

Our data in Figure 4 show that the defect in HA-M2 association in MDM is reversed upon changing residues 74-76, but not residues 71-73, in the M2 cytoplasmic tail to alanine. Of the three residues 74-76, previous studies showed that M2 with an amino acid substitution at E74 supports WT-level infectious virus release regardless of virus strains, whereas changes in E75 and Y76 reduce virus titer with the latter causing more severe defect (22). Y76 is further shown to be critical for vRNP incorporation and assembly/release of infectious IAV particles (22). To test whether the susceptibility to this suppression correlates with the ability of M2 to mediate vRNP incorporation into virions, we next analyzed M2 mutants with single point mutations at positions 74, 75 and 76 for HA-M2 association by PLA. As in Figure 4, only cells expressing M2 at the plasma membrane were analyzed for PLA between HA-M2. MDM infected with all three single mutant M2 viruses, E74A, E75A, and Y76A, showed abundant PLA signals between HA and M2 even under conditions where the actin cytoskeleton was not disrupted (Figures 5A and 5B). Upon actin disruption in MDM, no further increase in number of HA-M2 PLA spots was observed for the three M2 mutant viruses (Figures 5A and 5B). Of note, none of the mutations had any effect on HA-M2 association in dTHP1 cells as comparable number of PLA spots were observed on the surface of dTHP1 cells infected with the WT virus versus viruses encoding E74A, E75A, or Y76A M2 (Figures 5A and 5C).

**Figure 5:**
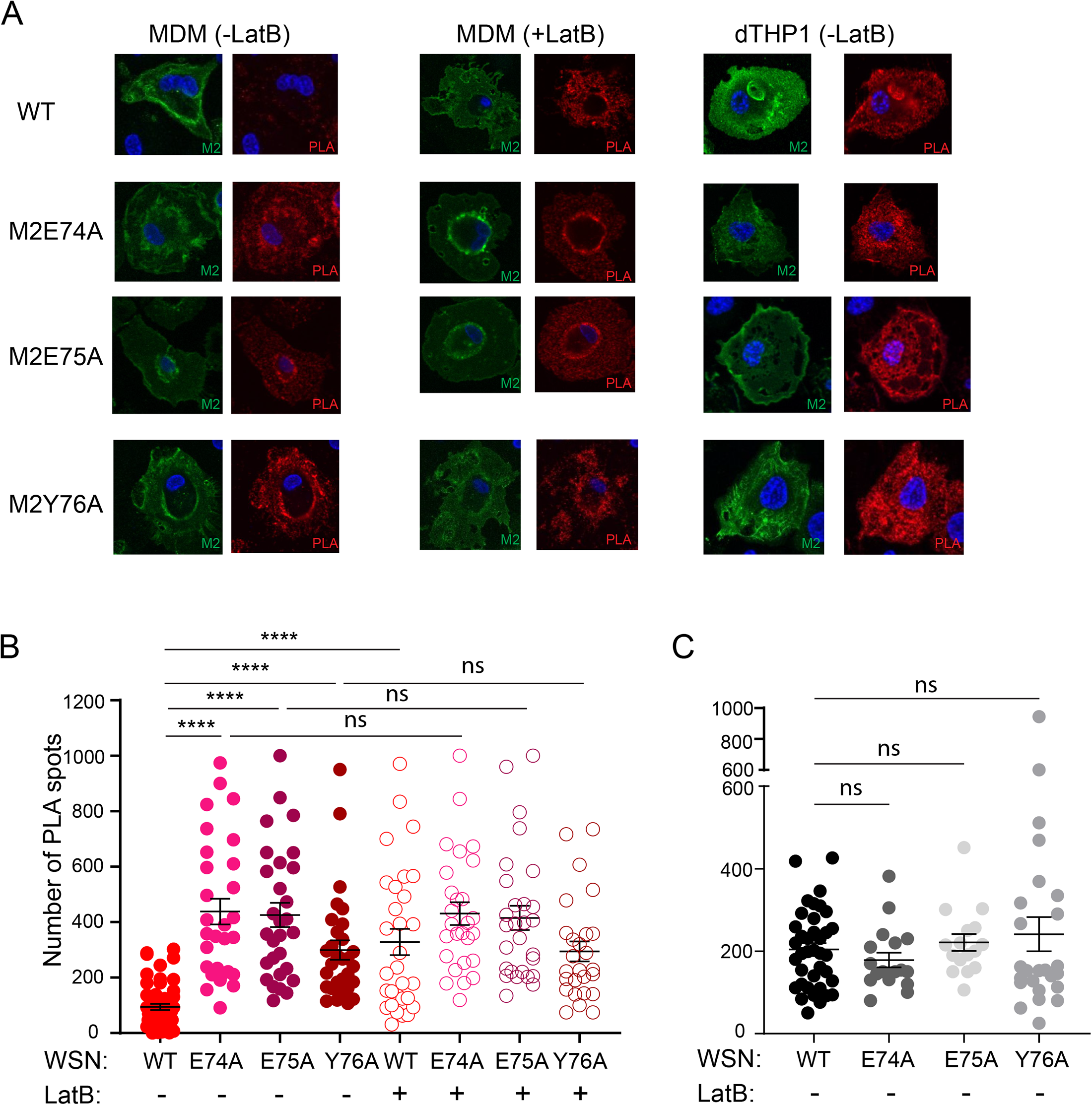
Amino acids at positions 74, 75, and 76 of the M2 protein are individually important for suppression of HA-M2 association in MDM. dTHP1 cells and MDM were infected with WT WSN or M2 mutant viruses at MOI 0.1 for 14 hours. Infected cells were treated with vehicle (DMSO) or 5 μM Lat B for 4 hours, before fixation. Fixed cells were examined by PLA using goat anti-HA and mouse anti-M2 antibodies. In addition, total surface M2 was also detected using a fluorescently-tagged anti-mouse IgG secondary antibody (shown in green). Nuclei were stained with DAPI (blue). (A) Representative maximum intensity projection images, which were reconstructed from z-stacks corresponding to the focal planes ranging from the middle plane of the nucleus to the bottom of the cells, are shown. (B,C) Number of PLA spots were counted for each cell. (B) Data are shown for experiments with MDM from three independent donors, and 8-10 cells were analyzed per experiment. (C) Data are shown for 2-3 experiments with dTHP1 cells and 8-10 cells were analyzed per experiment. Error bars represent standard error of mean. ****, p <0.0001; ns, non-significant.

Even though we specifically analyzed cells expressing M2 on the cell surface for PLA between HA and M2 in Figure 5, subtle differences in the cell surface expression between WT and mutant M2 could explain the differences in HA-M2 association in MDM. To address this possibility, we compared the cell surface expression of HA and M2 in MDM and dTHP1 cells infected with either WT or Y76A M2 virus using flow cytometry. Higher fractions of MDM were observed to be infected and hence expressing HA and M2 upon infection with the Y76A M2 virus than the WT virus (Figures 6A and 6B). Nevertheless, the mean fluorescence intensity (MFI), i.e., surface expression levels per cell, for the two viral proteins in positive cell populations was similar between WT and Y76A M2 viruses in both MDM and dTHP1 cells (Figure 6C). These results indicate that the plasma membrane trafficking of WT and Y76A M2 is equally efficient in dTHP1 cells and MDM and that enhanced association with HA observed for Y76A M2 relative to WT M2 is not due to enhanced surface expression of the M2 mutant.

**Figure 6:**
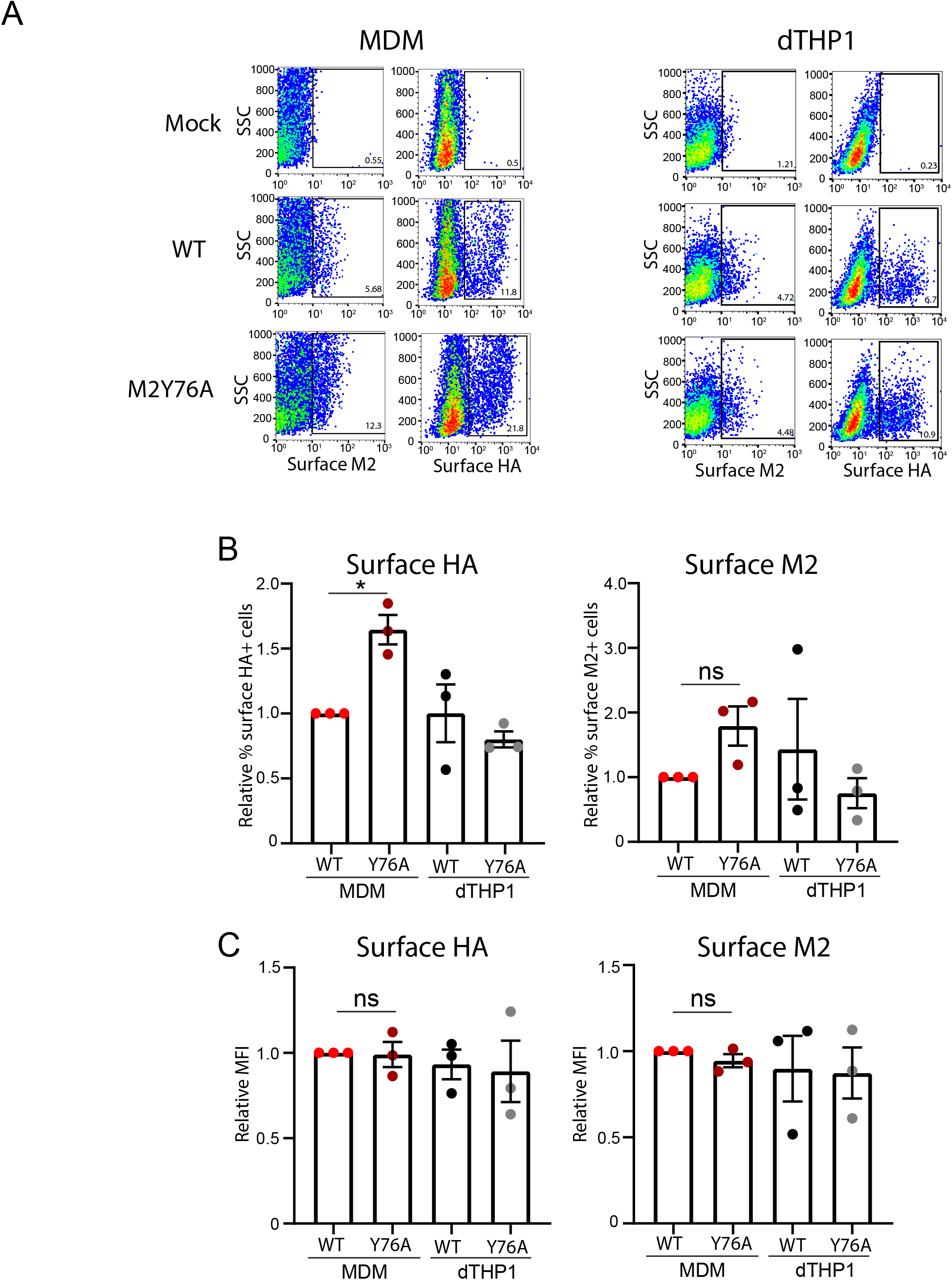
Cells infected WT and M2Y76A viruses express comparable levels of HA and M2 at the cell surface. dTHP1 cells and MDM were infected with WT WSN or the M2Y76A mutant virus at MOI 0.1. Infected cells were analyzed for cell surface expression of HA and M2 by flow cytometry at 18 hpi. (A) Representative flow plots for mock-infected (top row) and virus-infected (middle and bottom rows) cells are shown. Percentages of cells positive for viral proteins (boxed) are shown. (B) Relative percentages of cells positive for surface expression of HA (left) and M2 (right) are compared between WT and the Y76A mutant for both MDM and dTHP1 cells. (C) Relative MFIs for surface signal of indicated proteins for positive cell populations (gated in panel A) are shown. Data are shown as mean +/- SD and are from at least three independent experiments. *, p<0.05; ns, non-significant.

Overall, these data highlight the importance of the Glu-Glu-Tyr (EEY) sequence located in the cytoplasmic tail of M2 in MDM-specific inhibition of the association of M2 with HA. Notably, all three M2 mutants (E74A, E75A, Y76A) show the same phenotype in terms of reversing the defect in HA-M2 association in MDM. Since substitution of Y76 has previously been shown to diminish incorporation of vRNP into assembling virus particles, whereas substitution of E74 has no effect on infectious virus release (22), HA-M2 association is likely restricted in MDM in a manner independent of M2’s association with vRNP. Altogether, these results support the possibility that an Factin-dependent host factor expressed in MDM restricts the association of M2 with HA directly in a manner dependent on the EEY sequence in the M2 cytoplasmic domain rather than via other internal viral components.

## Discussion

In our previous study, we showed that both HA-M2 association and IAV assembly are suppressed by the actin cytoskeleton in MDM (15). In the current study, we sought to elucidate the molecular mechanism behind this defect in IAV assembly in MDM and to identify the viral factor under regulation in this cell type. Using a virus that fails to express M2 but expresses HA and M1 in infected cells (Figure 1), we showed that IAV assembly requires M2 expression in F-actin-disrupted MDM (Figure 2) and in dTHP1 cells (Figure 3). These data establish an important role for M2 during IAV particle assembly in human macrophages, which is consistent with the possibility in which MDM prevents recruitment of M2 to virus assembly sites enriched in HA, thereby suppressing IAV particle formation. We further showed that amino acid residues 74-76 in the cytoplasmic tail of M2 are individually important for the susceptibility of HA-M2 association to the MDM-specific inhibition (Figures 4 and 5), suggesting an M2-sequence-dependent mechanism behind restriction of HA-M2 association in this cell type. Overall, our study indicates that M2 is a viral factor that is essential for IAV assembly in MDM but also is a target of an MDM-specific regulation, which likely leads to the defect in HA-M2 association.

Previous studies using epithelial cells show that while M2 is required for pinching off of virus bud and its subsequent release, it is not required for induction of membrane curvature during particle formation (11, 13, 17, 23). It has also been proposed that M2 is recruited after induction of membrane curvature (41, 42), which is likely mediated by HA, NA or M1. At sites of virus assembly, M2 is enriched at the neck of the virus bud (17–19) and is thought to induce positive membrane curvature, which may be sufficient for membrane scission (17, 21). In contrast to epithelial cells, in both dTHP1 cells and MDM, M2 is essential for induction of membrane curvature during spherical IAV assembly (Figures 2 and 3). This difference between macrophages (dTHP1 and MDM) and epithelial cells may be due to the differences in the stoichiometry of other viral structural proteins at the plasma membrane of these different cell types. It is possible that in dTHP1 cells and MDM, other viral structural proteins, such as HA, NA, or M1, are either not present at the plasma membrane at a high enough concentration or do not cocluster to a high enough density, relative to epithelial cells. Under these circumstances, recruitment of M2 to HA-enriched virus budozones may be required to initiate particle assembly. Whether M2 directly initiates particle formation via its ability to alter membrane curvature or whether M2-mediated recruitment of M1 or vRNP to budozones induces particle formation remains to be determined.

The cytoplasmic tail of M2 plays an important role in the recruitment of M1 and/or vRNP to virus assembly sites. We found that the M2 region important for F-actin-dependent regulation of HA-M2 association in MDM (74-EEY-76) overlaps with the region that has been shown to function in recruitment of M1 and/or vRNP. However, the amino acid residue (E74) that has previously been shown to play little to no role in release of infectious virus particles (22) is still important for the susceptibility to inhibition of HA-M2 association in MDM. Moreover, the preceding amino acid sequence 71-SMR-73 in the M2 cytoplasmic tail is important for M1 recruitment (25) but does not play a role in regulation of HA-M2 association. Therefore, F-actin-dependent regulation of association of M2 with HA in MDM is likely to be independent of its association with M1 and/or vRNP. Considering that disruption of the EEY sequence relieves HA-M2 association from suppression by actin cytoskeleton, it is likely that MDM express a restriction factor that recognizes the EEY sequence. We favor a working hypothesis that an M2 EEY-binding protein inhibits HA-M2 association at the plasma membrane in a cytoskeleton-dependent manner, thereby preventing influenza A virus particle production in MDMs.

Overall, our study provides direct evidence for an M2-sequence-dependent mechanism for restriction of HA-M2 association in MDM. In addition, our data indicate that M2 plays an important role early during the assembly process in macrophages. Future comparative proteomic studies (43–46) between wild-type and mutant variants of M2 will potentially allow for identification of an MDM-specific host factor(s) that regulates HA-M2 association in an F-actin-dependent manner in MDM.

## Acknowledgements

We thank members of our laboratory for helpful discussions and critical reviews of the manuscript. This work is supported by NIH grant R21 AI 143276 (to A.O.).

## Notes

### Competing Interest Statement

The authors have declared no competing interest.

